# Dscam1 promotes blood cell survival in *Drosophila melanogaster* through a dual role in blood cells and neurons

**DOI:** 10.1101/2020.09.26.314997

**Authors:** Debra Ouyang, Xiaoyi Xiao, Anjeli Mase, Glenda Li, Sean Corcoran, Fei Wang, Katja Brückner

**Affiliations:** Eli and Edythe Broad Center of Regeneration Medicine and Stem Cell Research; Department of Cell and Tissue Biology; Cardiovascular Research Institute, University of California San Francisco, San Francisco, CA; Department of Molecular Biology, Princeton University, Princeton, NJ; Biological Science Research Center, Southwest University, Chongqing, China; Committee on Development, Regeneration, and Stem Cell Biology, The University of Chicago, IL

**Keywords:** Down Syndrome Cell Adhesion Molecule 1 (Dscam1), dreadlocks (dock), macrophage, hemocyte, sensory neuron, hematopoietic pocket, apoptosis, *Drosophila melanogaster*

## Abstract

Down Syndrome Cell Adhesion Molecule 1 (Dscam1) is a receptor-like cell adhesion molecule that is conserved across the animal kingdom, but its roles in hematopoiesis remain unknown. Dscam1 related genes in vertebrates and invertebrates are key regulators of neuron morphogenesis and neuronal tiling. In *Drosophila*, Dscam1 in addition has roles in blood cells (hemocytes) in innate immunity and phagocytosis of pathogens. Given the anatomical and functional role of peripheral sensory neurons as microenvironments for resident hematopoietic sites in the *Drosophila* larva, we sought to investigate the role of Dscam1 in this context. Interestingly, we find that Dscam1 fills the role of a previously anticipated factor in neuron-hemocyte communication that supports trophic survival: tissue specific silencing of *Dscam1* by in vivo RNAi in sensory neurons leads to neuron reduction, which in turn results in reduced hemocyte numbers due to apoptosis. Dscam1 silencing in hemocytes also results in a reduction of hemocytes and increased apoptosis. This cell-autonomous effect of *Dscam1* silencing can be mimicked by RNAi silencing of *dreadlocks* (*dock*), suggesting that intracellular Dscam1 signaling relies on the adapter protein Dock in this system. Our findings reveal a dual role for Dscam1 in *Drosophila* hematopoiesis, by promoting survival of the sensory neuron microenvironments that in turn support hemocyte survival, and by promoting survival of hemocytes cell-autonomously. It will be interesting to explore possible functions of vertebrate Dscam1 related genes such as DSCAML1 in blood cells and their trophic survival.

## 1. Introduction

Down Syndrome Cell Adhesion Molecule (Dscam) comprises a family of receptor-like cell adhesion molecules that are conserved across the animal kingdom (1, 2). In vertebrates, DSCAM and DSCAM-Like1 (DSCAML1) are known to play roles in nervous system development, including axon guidance, synaptic adhesion, and dendritic self-avoidance (3, 4). *Drosophila* Dscam1 is expressed in both neurons and blood cells (hemocytes) (5–7). In neurons, Dscam1 plays important roles in neuron guidance, dendrite formation, self avoidance of dendritic arbors and neuronal tiling of sensory neurons, i.e. the nonoverlapping coverage of neuronal dendrites from different but usually functionally related neurons (6, 8, 9) Dscam1 further has functional roles in hemocytes, specifically in the immune response including recognition and phagocytic uptake of bacteria (5, 10)

*Drosophila* Dscam1 comprises 10 immunoglobulin (Ig) domains, 6 fibronectin type III repeats, a single transmembrane domain, and a C-terminal cytoplasmic domain (9). *Drosophila* Dscam1 is expressed in many isoforms due to alternative splicing in exons 4,6,9 and 17 out of the total 24 exons (6); alternative splicing is estimated to enable expression of up to > 38,000 isoforms, or >18,000 isoforms in hemocytes (8, 11). Dscam1 engages in isoform specific homophilic binding and intracellular signaling (1, 9), e.g. adhesion signaling converting attachment to repulsion between sister neuronal branches, leading to dendrite repulsion (12). It further engages in heterotypic binding, such as with Netrins, and the Robo ligand Slit (13–15).

Hemocyte-specific loss of Dscam1 was reported to affect bacterial engulfment after infection (10). The bacterial binding capacity of Dscam1 isoforms was mapped to three out of ten Ig-like domains, and it was suggested that expression of 14–50 different isoforms of Dscam1 per hemocyte may provide an increased binding spectrum to pathogens (6, 16). However no adaptive responses in isoform variation could be detected following bacterial infection (17, 18).

Given the unique expression of Dscam1 in both neurons and hemocytes, we decided to study the role of Dscam1 in hematopoiesis, specifically in the paradigm of sensory neuron-regulated hematopoiesis in the *Drosophila* larva (Fig. 1 A). Here hemocytes of the embryonic lineage colonize segmentally repeated hematopoietic pockets (HPs) that contain sensory neuron microenvironments (19–22) (Fig. 1 B). This population of locally expanding hemocytes mainly consists of macrophage-like plasmatocytes, and parallels the independent myeloid system of self-renewing tissue macrophages in vertebrates (21–25). In the HPs, sensory neurons of the peripheral nervous system (PNS) promote hemocyte recruitment, survival, proliferation and transdifferentiation (19, 20). Previous work has shown that genetic ablation of PNS neurons affects the larval hemocyte pattern and leads to apoptosis, while induction of ectopic PNS neurons promotes hemocyte recruitment (20) Neuron-secreted Activin-β (Actβ) plays a role in the regulation of hemocyte proliferation and adhesion through Actβ pathway signaling (19). However, it remains unclear which neuron-produced cues may promote trophic survival of hemocytes, and whether membrane-bound adhesion receptors on hemocytes and/or sensory neurons may play a role in hematopoiesis in the HPs.

**Figure 1.**
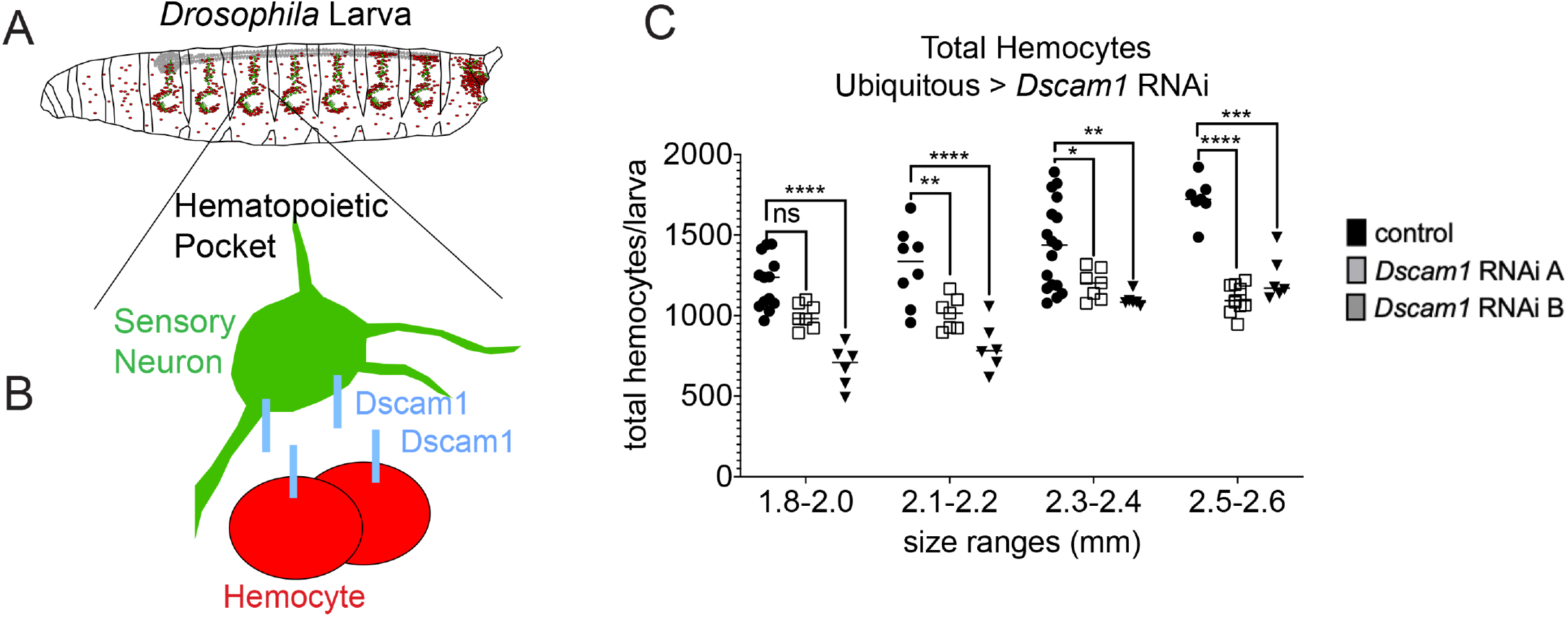
Dscam1 is expressed in both hemocytes and the sensory neuron microenvironment, and is needed to promote hemocyte numbers. (A) Model of *Drosophila* larva showing sensory neuron microenvironments and associated resident hemocytes. (B) Closeup of a hematopoietic pocket, illustrating Dscam1 expression in both sensory neurons and hemocytes. (C) Ubiquitous RNAi silencing of Dscam1 affects total hemocyte numbers in the *Drosophila* larva. Genotypes are control *ubi-GAL4, Hml*Δ-*DsRed/+* (n=45) and experiment *ubi-GAL4, Hml*Δ-*DsRed*/+; *UAS-Dscam1-RNAi* (line A Bloomington #29628 (n=32) or line B Bloomington #38542 (n=24))/+. Individual value plots with average and standard deviation; 2-way ANOVA, ns not significant; *,**,***,or **** corresponding to p≤0.05, 0.01, 0.001, or 0.0001.

In this study, we uncover a new dual role for Dscam1 in hematopoiesis of the embryonic blood cell lineage in the *Drosophila* larva. We show that Dscam1 has non-autonomous effects on hemocyte survival through its expression in sensory neurons, confirming their role as an essential microenvironment for hemocytes. We further reveal a cell-autonomous role of Dscam1 in hemocyte survival. Downstream of Dscam1, we find a role for the adapter protein dreadlocks (dock), suggesting that hemocyte survival signaling relies on the Dscam1-Dock axis.

## 2. Results

Intrigued by the dual expression of Dscam1 in hemocytes and sensory neurons in the Drosophila larval hematopoietic pockets (1, 5) (Fig. 1 A, B) we set out to investigate the functional role of Dscam1 in this system. We took an in vivo RNAi approach using the UAS/GAL4 system (35) as we intended to determine the role of Dscam1 at the ubiquitous level, and in distinct cell populations. First, we silenced Dscam1 ubiquitously using *ubi-GAL4* (29), and examined hemocytes, labeled by the transgenic fluorescent protein reporter *Hml*Δ-*DsRed* (20). Interestingly, total hemocytes were significantly reduced when *Dscam1* was silenced ubiquitously (Fig. 1 C). This observation was confirmed with two independent RNAi lines, suggesting that Dscam1 is required for hemocytes in the *Drosophila* larva.

Next we assessed the contribution of *Dscam1* expression in sensory neurons versus hemocytes. We silenced *Dscam1* in sensory neurons using the driver *21-7-GAL4* (20, 26) marked by coexpression of a *UAS-GFP* transgene; hemocytes were again visualized by expression of the transgenic reporter *Hml*Δ-*DsRed*. Comparing knockdowns by two alternative transgenic RNAi lines and control, we found that total hemocyte numbers per larva were significantly reduced upon *Dscam1* silencing in sensory neurons (Fig. 2 C). However, hemocyte localization in the segmentally repeated HPs remained fairly normal (Fig. 2 A, B).

**Figure 2.**
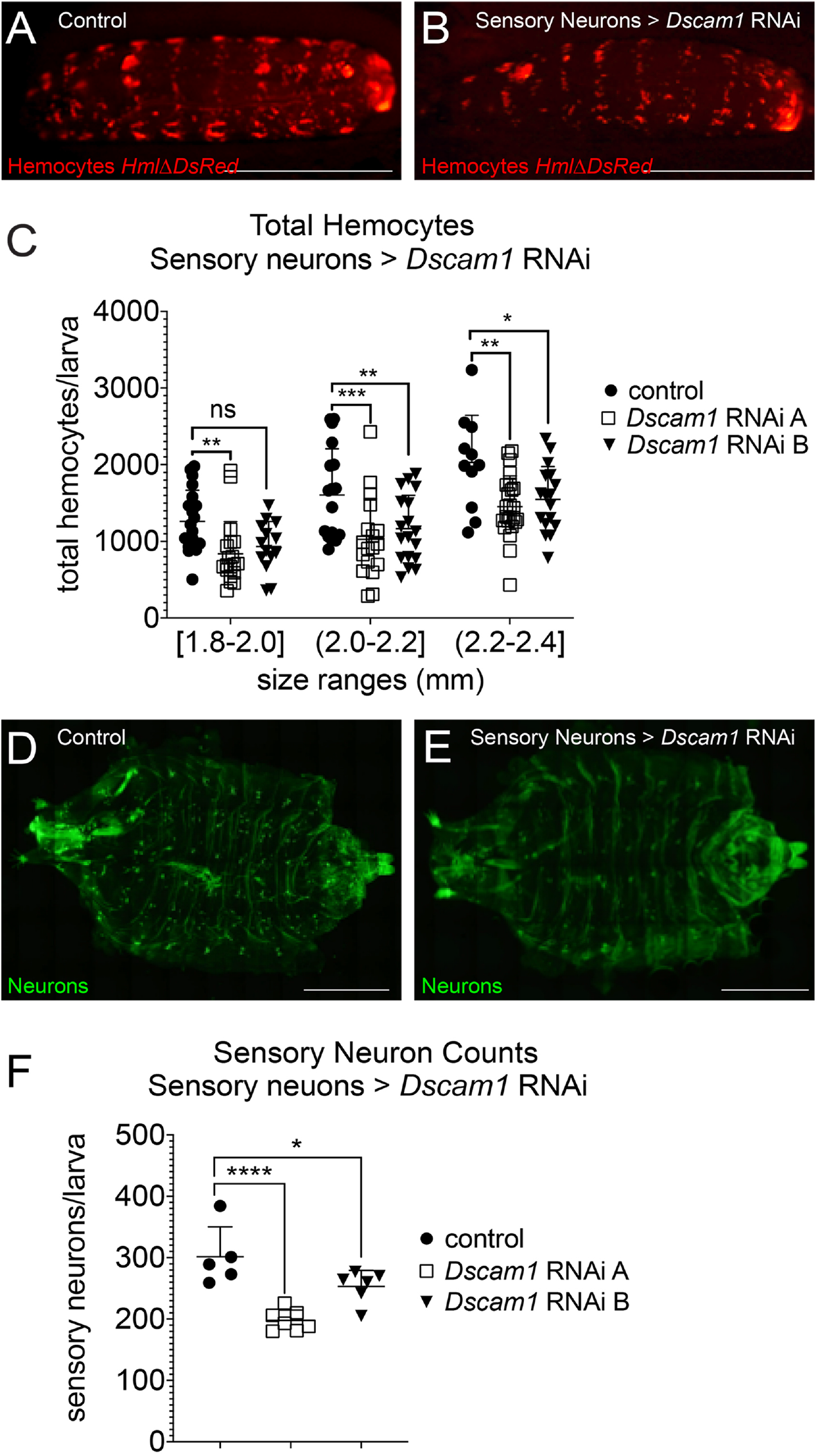
Effects of *Dscam1* silencing in sensory neurons: reduced neurons and reduced hemocytes. (A, B) Resident hemocyte pattern in *Drosophila* larvae, marked by *Hml*Δ-*DsRed* (red). (A) Control, genotype *21-7-GAL4, UAS-CD8-GFP, Hml*Δ-*DsRed*/+. (B) *Dscam1* knockdown, genotype *21-7-GAL4, UAS-CD8-GFP, Hml*Δ-*DsRed/+; UAS-Dscam1-RNAi* (line A Bloomington #29628 or line B Bloomington #38542)/+; scale bars 1mm. (C) RNAi silencing of *Dscam1* in sensory neurons affects total hemocyte numbers. Genotypes are control *21-7-GAL4, UAS-CD8-GFP* (n=49) and experiment *21-7-GAL4, UAS-CD8-GFP, Hml*Δ-*DsRed/+; UAS-Dscam1-RNAi* (line A Bloomington #29628 (n=61) or line B Bloomington #38542 (n=49))/+. Individual value plots with average and standard deviation; 2-way ANOVA, ns not significant; *, **, ***, or **** corresponding to p≤0.05, 0.01, 0.001, or 0.0001 (D, E) Larval fillets, hemocytes labeled by *21-7-GAL4* driving GFP in green. (D) Control genotype *21-7-GAL4, UAS-CD8-GFP, Hml*Δ-*DsRed*/+. (E) *Dscam1-RNAi* silencing in neurons genotype *21-7-GAL4, UAS-CD8-GFP, Hml*Δ-*DsRed/+; UAS-Dscam1-RNAi* (Bloomington #38542)/+. (F) Quantification of sensory neurons, genotypes are control *21-7-GAL4, UAS-CD8-GFP/+* (n=5) and *Dscam1* RNAi silencing in sensory neurons genotype *21-7-GAL4, UAS-CD8-GFP/+; UAS-Dscam1-RNAi* (line A Bloomington #29628 (n=6) or line B Bloomington #38542 (n=7))/+. Individual value plots with average and standard deviation; 1-way ANOVA, ns not significant; *, **, ***, or **** corresponding to p≤0.05, 0.01, 0.001, or 0.0001,

Questioning the basis of reduced hemocyte numbers upon *Dscam1* silencing in sensory neurons, we examined sensory neuron morphology and number. Strikingly, we found that sensory neurons were significantly reduced in *Dscam1* knockdowns compared to controls (Fig. 2 D-F). This situation recalls a previous report on the functional connection of hemocytes with sensory neurons, which showed that sensory neuron ablation in the larva, or *atonal* mutants that produce less sensory neurons, showed reduction of hemocytes due to apoptotic death (20). We conclude that Dscam1 is required in sensory neurons, and through an indirect, non-autonomous mechanism supports hemocytes.

Next we investigated the cell autonomous role of Dscam1 in hemocytes, silencing the gene using the double hemocyte driver line *Hml*Δ-*GAL4, UAS-GFP; He-GAL4* (27). Interestingly, we found that silencing of *Dscam1* in hemocytes also affected the total number of hemocytes per larva (Fig. 3 C); this phenotype was consistent among two independent RNAi lines. *Dscam1* silencing in hemocytes did not have a major effect on the resident hemocyte pattern (Fig. 3 A, B). However, we found a mildly increased but not statistically significant fraction of hemocytes with reduced adherence (‘bleed’ fraction of hemocytes) in younger animals, while this trend reverted in older larvae (Suppl. Fig. 1 A).

**Figure 3.**
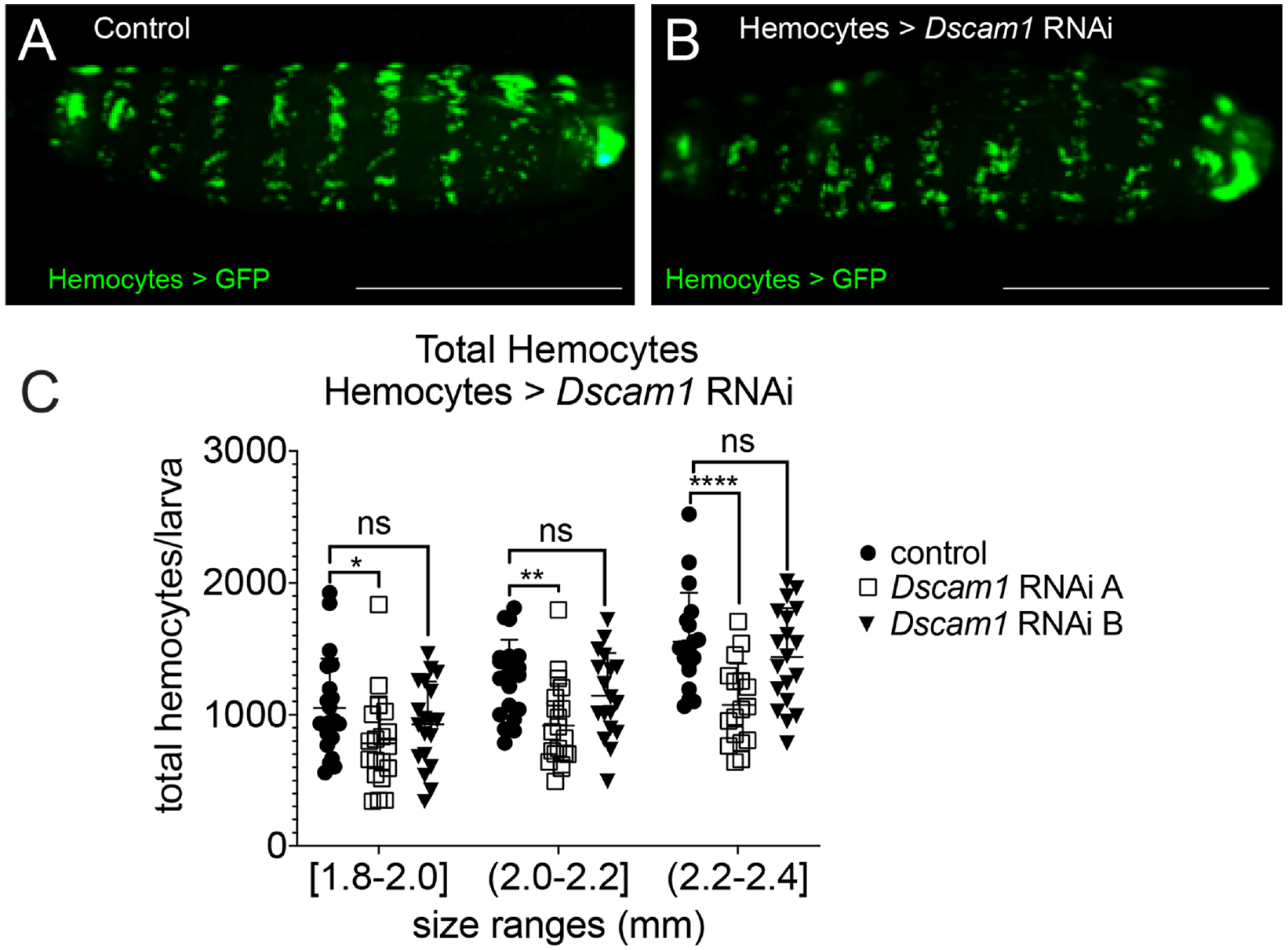
Cell-autonomous effects of *Dscam1* silencing in hemocytes. (A, B) Resident hemocyte pattern in *Drosophila* larvae, marked by GFP expression. (A) Control, genotype *Hml*Δ-*GAL4, UAS-GFP/+; He-GAL4*/+. (B) *Dscam1* knockdown, genotype *Hml*Δ-*GAL4, UAS-GFP/+; He-GAL4/UAS-Dscam1-RNAi* (Bloomington #29628); scale bars 1mm. (C) RNAi silencing of *Dscam1* in hemocytes affects total hemocyte numbers. Genotypes are experiment *Hml*Δ-*GAL4, UAS-GFP/+; He-GAL4/UAS-Dscam1-RNAi* (line A Bloomington #29628 (n=56) or line B Bloomington #38542 (n=56)) and control *Hml*Δ-*GAL4, UAS-GFP/+; He-GAL4/+*. (n=59). Individual value plots with average and standard deviation; 2-way ANOVA, ns not significant; *, **, ***, or **** corresponding to p≤0.05, 0.01, 0.001, or 0.0001.

Complementary to RNAi silencing, we asked whether *Dscam1* overexpression would have an effect on hemocyte numbers. Expressing *Dscam1* under control of the hemocyte driver *Hml*Δ-*GAL4; He-GAL4*, we found no phenotypic changes (Suppl. Fig. 1 B). We conclude that increased Dscam1 homophilic interactions among neighboring hemocytes do not seem to affect hemocytes. Overall the possibilities of overexpression experiments are limited though due to the lack of diverse Dscam1 isoforms in this approach, as typically unique Dscam1 isoforms determine specific interactions by homophilic pairings (7, 36).

In order to dissect the mechanism behind reduced hemocyte numbers, we determined hemocyte apoptosis. We assessed the fraction of apoptotic cells in *Dscam1* knockdowns and controls using NucView (Caspase-3 Substrate) labeling. Interestingly, *Dscam1* silencing in either hemocytes or sensory neurons resulted in an increase in the fraction of apoptotic hemocytes (Fig. 4 A, B). Our data suggest that Dscam1 is required both non-autonomously in sensory neurons, and cell autonomously in hemocytes, to promote hemocyte survival.

**Figure 4.**
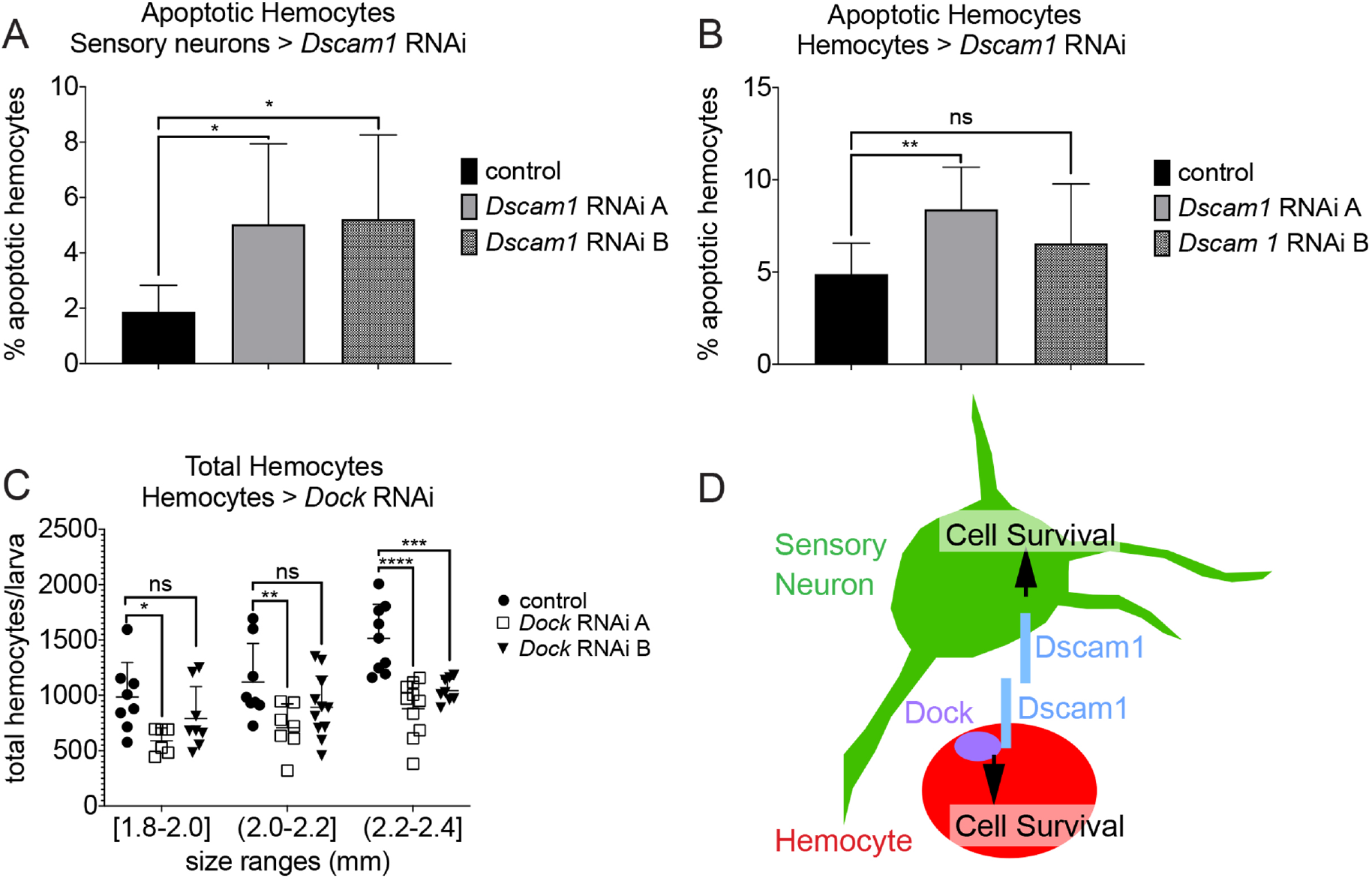
*Dscam1* silencing in sensory neurons or hemocytes results in hemocyte apoptosis; a role for Dock in hemocytes. (A, B) NucView labeling of apoptotic hemocytes. (A) *Dscam1* knockdown in sensory neurons, genotype *21-7-GAL4, UAS-CD8-GFP, Hml*Δ-*DsRed/+; UAS-Dscam1-RNAi* (line A Bloomington #29628 (n=10) or line B Bloomington #38542 (n=10))/+; and control genotype *21-7-GAL4, UAS-CD8-GFP, Hml*Δ-*DsRed/+* (n=9). (B) *Dscam1* knockdown in hemocytes, genotype *Hml*Δ-*GAL4, UAS-GFP/+; He-GAL4/ UAS-Dscam1-RNAi* (line A Bloomington #29628 (n=10) or line B Bloomington #38542 (n=10)); and control *Hml*Δ-*GAL4, UAS-GFP/+; He-GAL4/+* (n=10). Bar charts with average and standard deviation; 1-way ANOVA, ns not significant; *, **, ***, or **** corresponding to p≤0.05, 0.01, 0.001, or 0.0001. (C) RNAi silencing of *dock* in hemocytes affects total hemocyte numbers. Genotypes are experiment *Hml*Δ-*GAL4, UAS-GFP/+; He-GAL4/ UAS-dock-RNAi* (n=23) and control *Hml*Δ-*GAL4, UAS-GFP/+; He-GAL4*/+ (n=25). Individual value plots with average and standard deviation; 2-way ANOVA, ns not significant; *, **, ***, or **** corresponding to p≤0.05, 0.01, 0.001, or 0.0001. (D) Model. Dscam1 is required in both sensory neurons and hemocytes to promote hemocytes. In both cell types Dscam1 promotes cell survival; sensory neurons may provide unknown trophic cues to hemocytes, or the Dscam1 engagement itself may promote hemocyte survival. In hemocytes, signaling requires the adapter protein Dock that interacts with the Dscam1 cytoplasmic domain.

Lastly we asked whether Dscam1 in hemocytes might engage in intracellular signaling. We therefore tested the role of dreadlocks (dock), an SH2/SH3 domain adapter and major cytoplasmic interactor of the Dscam1 intracellular domain that has been involved in downstream signaling in neuronal models (6, 37). Interestingly, hemocyte-specific RNAi silencing of *dock* led to a reduction in hemocyte numbers, phenocopying, and in fact surpassing, the effects of *Dscam1* silencing (Fig. 4 C). This suggests that in *Drosophila* hemocytes, Dscam1 acts by intracellular signaling that involves the adapter protein Dock.

## 3. Discussion

Here we report a role of Dscam1 promoting blood cells of the hematopoietic pockets in the *Drosophila* larva. We find that Dscam1 supports hematopoiesis in a dual way; it is required in both sensory neurons that provide a microenvironment, and in hemocytes themselves, for proper blood cell survival, filling the role of a long-anticipated player that promotes hemocyte survival in this system (20). Dscam1 in sensory neurons acts non-autonomously on hemocytes, while it also promotes hemocytes in a cell-autonomous way (Fig. 4 D).

In sensory neurons, we find that *Dscam1* silencing results in reduction of these cells, suggesting that Dscam1 supports cell survival. Dscam1 is known for a variety of roles in the nervous system, such as dendritic self avoidance and mosaic tiling of sensory dendritic arborization neurons (12, 38–40) dendritic growth and branching in motor neurons (8), patterning of dendrites from olfactory projection neurons (41), and axon guidance (15, 42, 43).

It was demonstrated previously that hemocytes depend on the presence of sensory neurons, as animals with genetic ablation of sensory neurons, or *atonal* mutants that lack certain neurons, show increased apoptotic death of hemocytes and reduced total hemocyte numbers (20). Here, *Dscam1* knockdowns in sensory neurons, where a fraction of neurons is missing, phenocopy these hemocyte defects. These defects mainly comprise an increase in hemocyte apoptosis, while other aspects such as hemocyte adhesion seem less affected. We conclude that Dscam1 supports sensory neuron number and in turn promotes hemocyte survival. How sensory neurons communicate to hemocytes in support of their trophic survival remains subject of future study. One possibility is that sensory neurons produce an unrelated trophic survival factor that, when in limitation, starts leading to hemocyte death. Another interesting scenario is that homophilic binding of Dscam1 on sensory neurons to Dscam1 on hemocytes plays a role in sustaining trophic survival of hemocytes, while other aspects, such as hemocyte adhesion, are less affected. Dscam1 also has cell-autonomous functions in the hemocyte pool. *Dscam1* silencing causes reduction of hemocytes and increased apoptosis, suggesting that Dscam1 promotes or enables downstream trophic survival signaling. Surprisingly, Dscam1 silencing has no dramatic effects on hemocyte adhesion, with mild biphasic but mostly non-significant effects on hemocyte adhesion However, these effects are minimal compared to neuronal systems, suggesting a different role than in neuronal tiling, when Dscam1 on neighboring neuronal processes promotes repulsion (39, 40).

In our study we find that *Dscam1* overexpression in hemocytes does not show obvious phenotypes. This could be due to the fact that overexpression cannot mimic the splice variant diversity that Dscam1 naturally shows (7, 9), and that might be needed for effective Dscam1 neuron-hemocyte signaling. However in some cases, lack of Dscam1 diversity produces phenotypes, for example *Dscam1* overexpression was reported to impair precise neuronal connections (44). On the other hand, overexpression of one *Dscam1* splice variant may enable heterotypic interactions as they were reported e.g. with Netrins, the Robo1 ligand Slit, and Robo1 itself (13–15). Moreover, in our system, homophilic interactions would be expected to take place between neighboring, *Dscam1*-overepressing hemocytes themselves. Since we find that Dscam1 promotes hemocyte survival, it is possible though that *Dscam1* overexpression may not cause an obvious phenotype, as it does not promote hemocyte survival beyond the already high levels of survival under control conditions.

How does Dscam1 act cell-autonomously in hemocytes? Several signaling mechanisms have been described for Dscam1, from cytoplasmic domain signaling interactors (6, 37) to the cleavage and nuclear translocation of the Dscam1 intracellular domain, which resembles Notch signaling (45). We find that the adaptor protein Dock is required to maintain hemocyte numbers. Dock is a SH2/SH3 domain adapter homologous to human Nck (46); both have been implicated in tyrosine kinase signaling (46, 47). Dock is required for Dscam1 functions in the nervous system such as axon guidance and dendritogenesis (6, 48), through direct interaction with the Dscam1 cytoplasmic domain (6, 37, 48). Dock in turn binds and activates the p21-activated kinase Pak (37, 49, 50), an effector of the Rho GTPases Cdc42 and Rac (46, 48, 49, 51)

Our work reveals an unexpected role of Dscam1/Dock signaling in promoting anti-apoptotic survival of *Drosophila* hemocytes. This activity has not been described previously, although it has e.g. been reported that loss of the Dock homolog Nck increases UV-induced p53 phosphorylation and apoptosis in human cells (52). Moreover, the downstream kinase Pak1 phosphorylates the proapoptotic protein bad and protects murine cells from apoptosis (53), and Pak1 is also involved in prevention of apoptosis in Xenopus oocytes (54). Interestingly, our data indicate that *dock* silencing may have a stronger phenotype than *Dscam1* silencing, suggesting that Dock may act downstream of additional signaling players in hemocytes, which will be subject to future study. Likewise, it will be interesting to determine the role of Pak and other signaling components in hemocyte survival.

The *Drosophila* Dscam1-related vertebrate DSCAM and DSCAML1 are well known for their roles in neuronal development (3, 4). Interestingly, DSCAML1 is expressed in various cell types of the immune system, such as T- and B-lymphocytes and other cells of peripheral blood and bone marrow (The Human Protein Atlas). Our findings raise the question whether vertebrate DSCAML1 may also have functions in vertebrate hematopoiesis, in particular in the trophic survival of immune cells, and possibly their interactions with local neurons or sensors that may provide hematopoietic microenvironments.

## 4. Experimental Procedures

### Drosophila Strains

*Drosophila* lines used were *Hml*Δ-*DsRed* (20); *21-7-GAL4* (20, 26); *Hml*Δ-*GAL4, UAS-GFP; He-GAL4* (kind gift of Jesper Kronhamn and Dan Hultmark (27); *Hml*Δ-*GAL4* (28)), *ubi-GAL4/CyO* (Bloomington BL# 32551 (29)) *UAS-Dscam1-RNAi* (Bloomington TRiP, BL#29628), *UAS-Dscam1-RNAi* (Bloomington, TRiP BL#38945). *UAS-Dscam1* (Bloomington BL#66202); *UAS-Dscam1* (BL#66203); *UAS-dock-RNAi*; ((Bloomington TRiP, BL#27728), *yw* (from Perrimon lab). To express transgenes in PNS sensory neurons and visualize hemocytes, we used the PNS driver *21-7 GAL4* marked by expression of *UAS CD8 GFP* (30) combined with the hemocyte reporter *Hml*Δ-*DsRed* (20). To express transgenes in hemocytes and visualize hemocytes, we used the hemocyte driver *Hml*Δ-*GAL4, UAS-GFP; He-GAL4* which also marks hemocytes by the co-expressed *UAS-GFP*. Both lines used for visualization of hemocytes were crossed to *UAS-RNAi* lines or control (*yw*). To visualize and count sensory neurons, we used *21-7-GAL4, UAS CD8 GFP*, crossed to *RNAi* lines or control (*yw*). Fly crosses were maintained at 25°C.

### Hemocyte Releases and Quantification

Hemocyte releases were performed according to (31, 32), determining total hemocyte numbers, or separately quantifying circulating and resident hemocytes as ‘bleed’ and ‘scrape’ fractions, respectively. The latter were used to calculate the fraction of circulating hemocytes (32). Larvae of 2^nd^ instar stage were analyzed at size ranges corresponding to hours after egg laying (AEL): 1.8-2.0mm (54-59h AEL), 2.01-2.2mm (60-65h AEL), and 2.21-2.4mm (66-71h AEL). Hemocytes were released onto glass slides with wells marked by a PAP pen (Beckman Coulter) or in multi well cell culture dishes. Tile scan images of wells with released hemocytes were taken on a Leica DMI4000B fluorescence microscope with Leica DFC350FX camera at 20x magnification. Hemocyte counts were determined by semi-automated quantification using Metamorph or ImageJ/ Fiji (33) software (32). Larvae were not separated by gender, assuming equal fractions of male and female larvae. For hemocyte numbers of each genotype and condition, the mean and standard deviation were determined and significance was tested by 2-way ANOVA (Prism).

### Larval Fillets and Neuron Quantification

*Drosophila* larval fillets were prepared according to (20, 34). Because of size constraints for dissections, 3^rd^ instar larvae were examined. Larvae were pinned on silicone plates filled with PBS and cut open dorsally using iris scissors (Fine Science Tools). Organs such as intestine, fat body, and tracheal tubes were removed, while extra care was taken to keep the brain intact to the larval body. The tissue was fixed with 4% paraformaldehyde in PBS, washed and mounted with a cover slip on a glass slide. Imaging was done on a Leica DMI4000B fluorescence microscope. Tile scan images were taken at 20x magnification and sensory neurons of the body wall were counted manually. Average, standard deviation and 1-way ANOVA were calculated (Prism).

### Apoptosis Assay

Hemocytes from larvae were released into wells marked by a hydrophobic PAP pen (Beckman Coulter) on glass slides, filled with 20-30 μL PBS. NucView Caspase-3 substrate detection (green or red label to complement fluorescent hemocyte reporters) was performed according to the manufacturer’s instructions (Biotium); cells were incubated for 1 hour at room temperature. Cells were imaged by fluorescence tile scan microscopy on Leica DMI4000B microscope with Leica DFC350FX camera and 20x objective. Fractions of NucView positive hemocytes were determined, average and standard deviations were calculated, and significance of changes across genotypes was determined by 1-way ANOVA (Prism).

### Microscopy and Imaging

Live fluorescence imaging of larvae expressing GFP and/or DsRed was performed as described previously (20). 2^nd^ and early 3^rd^ instar larvae corresponding to stages analyzed in total hemocyte releases were imaged. Imaging was performed using a Leica M205FA fluorescence stereoscope with DFC300FX color digital camera and Leica LAS Montage module, combining z-stacks into single in-focus images.

## Authors’ contributions

KB conceived and supervised the study. DO, XX, AM, GL carried out experiments; SC generated a key *Drosophila* line. DO, XX, AM, SC and KB analyzed the data. SC and AM prepared files and figures. KB wrote the manuscript, with input from all authors.

## Acknowledgements

We thank J. Kronhamn and D. Hultmark, Y.N. Jan, N. Perrimon, TRiP, and the Bloomington Stock Center for fly stocks. Thanks to members of the Brückner lab for feedback on the project and manuscript.

---

**Supplemental Figure 1.**
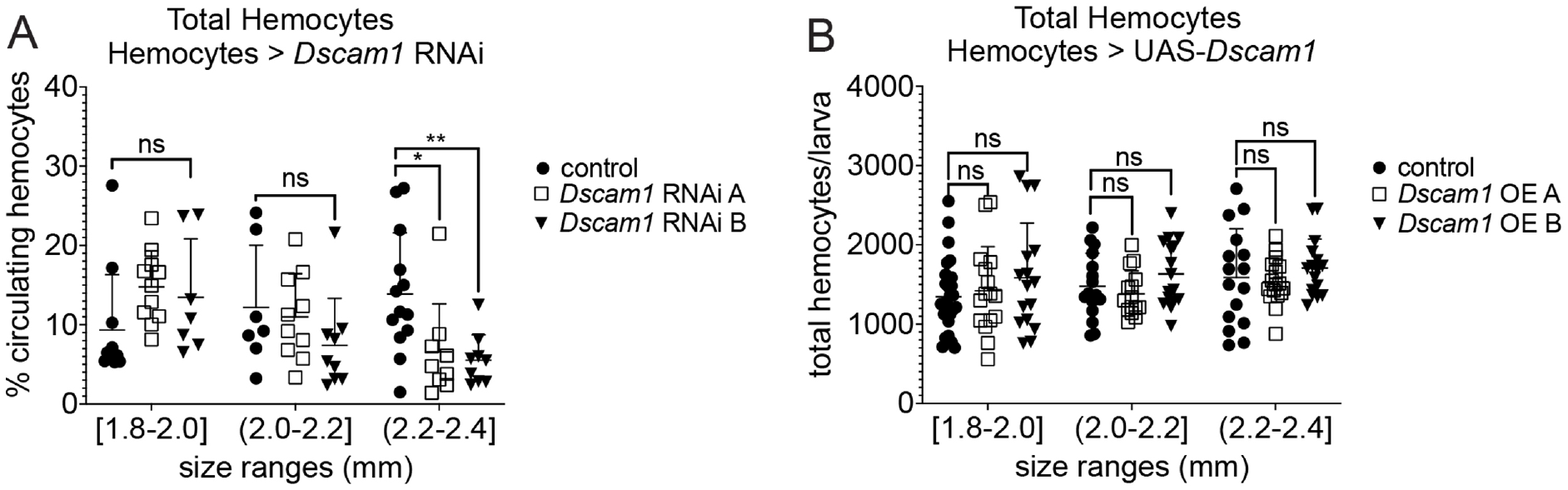
Effect of *Dscam1* silencing on hemocyte residence; *Dscam1* overexpression does not affect total hemocytes. (A) RNAi silencing of *Dscam1* in hemocytes has only minor effects of hemocyte residence. Percentage of circulating hemocytes was determined. Genotypes are experiment *Hml*Δ-*GAL4, UAS-GFP/+; He-GAL4/ UAS-Dscam1-RNAi* (line A Bloomington #29628 (n=30) or line B Bloomington #38542 (n=25)) and control *Hml*Δ-*GAL4, UAS-GFP/+; He-GAL4/+* (n=31). Individual value plots with average and standard deviation; 2-way ANOVA, ns not significant; *, **, ***, or **** corresponding to p≤0.05, 0.01, 0.001, or 0.0001. (B) Overexpression of *Dscam1* in hemocytes has no significant effect on total hemocytes. Genotypes are experiment *Hml*Δ-*GAL4, UAS-GFP/+; He-GAL4/ UAS-Dscam1* (*UAS-Dscam1* OE line A Bloomington BL#66202 (n=48) or OE line B BL#66203 (n=48)) and control *Hml*Δ-*GAL4, UAS-GFP/+; He-GAL4/+* (n=60). Individual value plots with average and standard deviation; 2-way ANOVA, ns not significant; *, **, ***, or **** corresponding to p≤0.05, 0.01, 0.001, or 0.0001.

## References

1. Schmucker D, Chen B. Dscam and DSCAM: complex genes in simple animals, complex animals yet simple genes. Genes Dev. 2009;23(2):147–56.

2. Armitage SA, Freiburg RY, Kurtz J, Bravo IG. The evolution of Dscam genes across the arthropods. BMC Evol Biol. 2012;12:53.

3. Montesinos ML. Roles for DSCAM and DSCAML1 in central nervous system development and disease. Adv Neurobiol. 2014;8:249–70.

4. Barlow GM, Micales B, Chen XN, Lyons GE, Korenberg JR. Mammalian DSCAMs: roles in the development of the spinal cord, cortex, and cerebellum? Biochem Biophys Res Commun. 2002;293(3):881–91.

5. Watson FL, Puttmann-Holgado R, Thomas F, Lamar DL, Hughes M, Kondo M, et al. Extensive diversity of Ig-superfamily proteins in the immune system of insects. Science. 2005;309(5742):1874–8.

6. Schmucker D, Clemens JC, Shu H, Worby CA, Xiao J, Muda M, et al. *Drosophila* Dscam is an axon guidance receptor exhibiting extraordinary molecular diversity. Cell. 2000;101(6):671–84.

7. Schmucker D, Flanagan JG. Generation of recognition diversity in the nervous system. Neuron. 2004;44(2):219–22.

8. Hutchinson KM, Vonhoff F, Duch C. Dscam1 is required for normal dendrite growth and branching but not for dendritic spacing in *Drosophila* motoneurons. J Neurosci. 2014;34(5):1924–31.

9. Wojtowicz WM, Flanagan JJ, Millard SS, Zipursky SL, Clemens JC. Alternative splicing of *Drosophila* Dscam generates axon guidance receptors that exhibit isoform-specific homophilic binding. Cell. 2004;118(5):619–33.

10. Vlisidou I, Wood W. *Drosophila* blood cells and their role in immune responses. Febs J. 2015;282(8):1368–82.

11. Zipursky SL, Wojtowicz WM, Hattori D. Got diversity? Wiring the fly brain with Dscam. Trends Biochem Sci. 2006;31(10):581–8.

12. Matthews BJ, Kim ME, Flanagan JJ, Hattori D, Clemens JC, Zipursky SL, et al. Dendrite self-avoidance is controlled by Dscam. Cell. 2007;129(3):593–604.

13. Ly A, Nikolaev A, Suresh G, Zheng Y, Tessier-Lavigne M, Stein E. DSCAM is a netrin receptor that collaborates with DCC in mediating turning responses to netrin-1. Cell. 2008;133(7):1241–54.

14. Matthews BJ, Grueber WB. Dscam1-mediated self-avoidance counters netrin-dependent targeting of dendrites in *Drosophila*. Curr Biol. 2011;21(17):1480–7.

15. Alavi M, Song M, King GL, Gillis T, Propst R, Lamanuzzi M, et al. Dscam1 Forms a Complex with Robo1 and the N-Terminal Fragment of Slit to Promote the Growth of Longitudinal Axons. PLoS Biol. 2016;14(9):e1002560.

16. Wojtowicz WM, Wu W, Andre I, Qian B, Baker D, Zipursky SL. A vast repertoire of Dscam binding specificities arises from modular interactions of variable Ig domains. Cell. 2007;130(6):1134–45.

17. Armitage SA, Sun W, You X, Kurtz J, Schmucker D, Chen W. Quantitative profiling of *Drosophila melanogaster* Dscam1 isoforms reveals no changes in splicing after bacterial exposure. PLoS One. 2014;9(10):e108660.

18. Peuss R, Wensing KU, Woestmann L, Eggert H, Milutinovic B, Sroka MG, et al. Down syndrome cell adhesion molecule 1: testing for a role in insect immunity, behaviour and reproduction. R Soc Open Sci. 2016;3(4):160138.

19. Makhijani K, Alexander B, Rao D, Petraki S, Herboso L, Kukar K, et al. Regulation of *Drosophila* hematopoietic sites by Activin-beta from active sensory neurons. Nat Commun. 2017;8:15990.

20. Makhijani K, Alexander B, Tanaka T, Rulifson E, Brückner K. The peripheral nervous system supports blood cell homing and survival in the *Drosophila* larva. Development. 2011;138:5379–91.

21. Gold KS, Brückner K. *Drosophila* as a model for the two myeloid blood cell systems in vertebrates. Exp Hematol. 2014;42(8):717–27.

22. Gold KS, Brückner K. Macrophages and cellular immunity in *Drosophila melanogaster*. Semin Immunol. 2015;27(6):357–68.

23. Davies LC, Jenkins SJ, Allen JE, Taylor PR. Tissue-resident macrophages. Nat Immunol. 2013;14(10):986–95.

24. Perdiguero EG, Klapproth K, Schulz C, Busch K, Azzoni E, Crozet L, et al. Tissue-resident macrophages originate from yolk-sac-derived erythro-myeloid progenitors. Nature. 2014.

25. Sieweke MH, Allen JE. Beyond stem cells: self-renewal of differentiated macrophages. Science. 2013;342(6161):1242974.

26. Song W, Onishi M, Jan LY, Jan YN. Peripheral multidendritic sensory neurons are necessary for rhythmic locomotion behavior in *Drosophila* larvae. Proc Natl Acad Sci U S A. 2007;104(12):5199–204.

27. Yang H, Kronhamn J, Ekstrom JO, Korkut GG, Hultmark D. JAK/STAT signaling in *Drosophila* muscles controls the cellular immune response against parasitoid infection. EMBO Rep. 2015;16(12):1664–72.

28. Sinenko SA, Mathey-Prevot B. Increased expression of *Drosophila* tetraspanin, Tsp68C, suppresses the abnormal proliferation of ytr-deficient and Ras/Raf-activated hemocytes. Oncogene. 2004;23(56):9120–8.

29. Schulz C, Perezgasga L, Fuller MT. Genetic analysis of dPsa, the *Drosophila* orthologue of puromycin-sensitive aminopeptidase, suggests redundancy of aminopeptidases. Dev Genes Evol. 2001;211(12):581–8.

30. Lee T, Luo L. Mosaic analysis with a repressible cell marker for studies of gene function in neuronal morphogenesis. Neuron. 1999;22(3):451–61.

31. Corcoran S, Brückner K. Quantification of blood cells in *Drosophila* and other insects. Springer Protocols Handbooks: Immunity in Insects. 2020;Sandrelli F, Tettamanti G, editors:65–77.

32. Petraki S, Alexander B, Brückner K. Assaying Blood Cell Populations of the *Drosophila melanogaster* Larva. J Vis Exp. 2015(105).

33. Schindelin J, Arganda-Carreras I, Frise E, Kaynig V, Longair M, Pietzsch T, et al. Fiji: an open-source platform for biological-image analysis. Nat Methods. 2012;9(7):676–82.

34. Tenenbaum CM, Gavis ER. Removal of *Drosophila* Muscle Tissue from Larval Fillets for Immunofluorescence Analysis of Sensory Neurons and Epidermal Cells. J Vis Exp. 2016(117).

35. Brand AH, Perrimon N. Targeted gene expression as a means of altering cell fates and generating dominant phenotypes. Development. 1993;118(2):401–15.

36. Hattori D, Millard SS, Wojtowicz WM, Zipursky SL. Dscam-mediated cell recognition regulates neural circuit formation. Annu Rev Cell Dev Biol. 2008;24:597–620.

37. Li W, Guan KL. The Down syndrome cell adhesion molecule (DSCAM) interacts with and activates Pak. J Biol Chem. 2004;279(31):32824–31.

38. Hughes ME, Bortnick R, Tsubouchi A, Baumer P, Kondo M, Uemura T, et al. Homophilic Dscam interactions control complex dendrite morphogenesis. Neuron. 2007;54(3):417–27.

39. Soba P, Zhu S, Emoto K, Younger S, Yang SJ, Yu HH, et al. *Drosophila* sensory neurons require Dscam for dendritic self-avoidance and proper dendritic field organization. Neuron. 2007;54(3):403–16.

40. Grueber WB, Jan LY, Jan YN. Tiling of the *Drosophila* epidermis by multidendritic sensory neurons. Development. 2002;129(12):2867–78.

41. Zhu H, Hummel T, Clemens JC, Berdnik D, Zipursky SL, Luo L. Dendritic patterning by Dscam and synaptic partner matching in the *Drosophila* antennal lobe. Nat Neurosci. 2006;9(3):349–55.

42. Zhan XL, Clemens JC, Neves G, Hattori D, Flanagan JJ, Hummel T, et al. Analysis of Dscam diversity in regulating axon guidance in *Drosophila* mushroom bodies. Neuron. 2004;43(5):673–86.

43. Andrews GL, Tanglao S, Farmer WT, Morin S, Brotman S, Berberoglu MA, et al. Dscam guides embryonic axons by Netrin-dependent and - independent functions. Development. 2008;135(23):3839–48.

44. Cvetkovska V, Hibbert AD, Emran F, Chen BE. Overexpression of Down syndrome cell adhesion molecule impairs precise synaptic targeting. Nat Neurosci. 2013;16(6):677–82.

45. Sachse SM, Lievens S, Ribeiro LF, Dascenco D, Masschaele D, Horre K, et al. Nuclear import of the DSCAM-cytoplasmic domain drives signaling capable of inhibiting synapse formation. Embo J. 2019;38(6).

46. Li W, Fan J, Woodley DT. Nck/Dock: an adapter between cell surface receptors and the actin cytoskeleton. Oncogene. 2001;20(44):6403–17.

47. Garrity PA, Rao Y, Salecker I, McGlade J, Pawson T, Zipursky SL. *Drosophila* photoreceptor axon guidance and targeting requires the dreadlocks SH2/SH3 adapter protein. Cell. 1996;85(5):639–50.

48. Kamiyama D, McGorty R, Kamiyama R, Kim MD, Chiba A, Huang B. Specification of Dendritogenesis Site in *Drosophila* aCC Motoneuron by Membrane Enrichment of Pak1 through Dscam1. Dev Cell. 2015;35(1):93–106.

49. Hing H, Xiao J, Harden N, Lim L, Zipursky SL. Pak functions downstream of Dock to regulate photoreceptor axon guidance in *Drosophila*. Cell. 1999;97(7):853–63.

50. Perez-Nunez R, Barraza N, Gonzalez-Jamett A, Cardenas AM, Barnier JV, Caviedes P. Overexpressed Down Syndrome Cell Adhesion Molecule (DSCAM) Deregulates P21-Activated Kinase (PAK) Activity in an In Vitro Neuronal Model of Down Syndrome: Consequences on Cell Process Formation and Extension. Neurotox Res. 2016;30(1):76–87.

51. Rane CK, Minden A. P21 activated kinases: structure, regulation, and functions. Small GTPases. 2014;5.

52. Errington TM, Macara IG. Depletion of the adaptor protein NCK increases UV-induced p53 phosphorylation and promotes apoptosis. PLoS One. 2013;8(9):e76204.

53. Schurmann A, Mooney AF, Sanders LC, Sells MA, Wang HG, Reed JC, et al. p21-activated kinase 1 phosphorylates the death agonist bad and protects cells from apoptosis. Mol Cell Biol. 2000;20(2):453–61.

54. Faure S, Vigneron S, Doree M, Morin N. A member of the Ste20/PAK family of protein kinases is involved in both arrest of Xenopus oocytes at G2/prophase of the first meiotic cell cycle and in prevention of apoptosis. Embo J. 1997;16(18):5550–61.

